# The fungicide Tebuconazole induces electromechanical cardiotoxicity in murine hearts

**DOI:** 10.1101/2021.10.13.464222

**Authors:** Artur Santos-Miranda, Julliane V. Joviano-Santos, Taynara Cruz-Nascimento, Diego Santos Souza, Leisiane Marques, Danilo Roman-Campos

## Abstract

Tebuconazole (TEB) is an important fungicide that belongs to the triazole family. It is largely applied in agriculture and its use has increased in the last decade. Since TEB is stable in water and soil, long-term exposure of humans to this pesticide is a real threat. Acute toxicological studies to uncover the toxicity of TEB are limited, and there is evidence of an association between long-term exposure to TEB and damage of several biological systems, including hepatotoxicity and cardiotoxicity. In this paper, the effects of acute exposure of cardiomyocytes and murine hearts to TEB were addressed to elucidate its impact on electromechanical properties of the cardiac tissue. In whole-cell patch-clamp records, TEB inhibited both the total outward potassium current (IC_50_=5.7±1.5 μmol.l^-1^) and the L-type calcium current (IC_50_=33.2±7.4 μmol.l^-1^). Acute exposure to TEB at 30 μmol.l^-1^ prolonged the action potential duration as well as an induced out-of-pace action potential, and increased the sodium/calcium exchanger current in its forward and reverse modes. Moreover, sarcomere shortening and calcium transient in isolated cardiomyocytes was enhanced when cells were exposed to TEB at 30 μmol.l^-1^. In *ex vivo* experiments, TEB 30 μmol.l^-1^ caused significant electrocardiogram remodeling with prolonged PR, QRS, and QT interval duration. Accordingly, TEB exposure was prone to the appearance of arrhythmias. Combined, our results demonstrate that acute TEB exposure affects the cardiomyocyte’s electro-contractile properties and triggers the appearance of ECG abnormalities, including conduction defects and arrhythmias.

## Introduction

Tebuconazole (TEB) [(RS)-1-p-chlorophenyl-4,4-dimethyl-3-(1H-1,2,4-triazol-1-yl methyl)pentan-3-ol] is a large-spectrum fungicide with plant growth-regulating properties that belongs to the triazole group. Today, triazoles correspond to 30% of marketed fungicides (Shishatskaya et al., 2018), and their mechanism of action is attributed to inhibition of the lanosterol-14-α-demethylase activity, which is a cytochrome P450 enzyme (Kwok and Loeffler, 1993; Podust et al., 2001; Yuzo and Yuri, 1987). Given its high fungicide effectiveness, TEB use has consistently grown, becoming a sales leader (Bowen et al., 1997; Toda et al., 2021). Only in the United States, TEB use has increased from less than 0.1 million pounds in 1994 to more than 2 million pounds in 2017 (National Water-Quality Assessment Project, 2017). The widespread use of TEB combined with its physical-chemical stability (low solubility and no photolysis in water) (Lewis et al., 2016; U. S. EPA, 2000), lead to frequent detection of the fungicide in food and water reservoirs around the globe (Chau et al., 2015; Kahle et al., 2008; Montuelle et al., 2010; Rabiet et al., 2010; Richardson, 2009). Thus, human exposure to TEB may occur due to its presence in food or drinking water, and also through inhalation or dermal contact in rural areas where this fungicide is broadly applied (Mercadante et al., 2014; Schummer et al., 2012; U.S. EPA, 2021). Indeed, TEB-OH and TEB-COOH, the urine metabolites of TEB, have been detected in urine samples from agricultural workers (Fustinoni et al., 2012). Therefore, despite the benefits of TEB for agriculture, it has potential implications for human health.

Experimental evidence suggests that TEB may have harmful effects on nontarget organisms such as humans (Zhou et al., 2016), fishes (Castro et al., 2018; Li et al., 2019), and rodents (EFSA, 2014). Long-term exposition to the pesticide has been implicated to induce carcinogen (Hu et al., 2007; U.S. EPA, 2010), and liver dysfunction in mice (Ku et al., 2021). TEB also causes impairment of neurological, immunological, and reproductive functions in adult rats (Moser et al., 2001). Consistent with other triazole fungicides, TEB has endocrine-disrupting effects (Taxvig et al., 2007), alters the cytochrome P450 enzymes activity, and causes oxidative stress in the rat liver (Martin et al., 2007; Yang et al., 2018). However, toxicological evaluation covering its potential cardiotoxicity is much less explored. Some recent studies described that TEB may induce heart injury (Ben Othmène et al., 2020a), with myocardial fibrosis. Oral administration of TEB during 28 days induced oxidative stress and histopathological alterations in rats’ hearts (Ben Othmène et al., 2020b). Moreover, TEB changed lipid profile, induced cell apoptosis and DNA damage (Ben Othmène et al., 2020c).

Despite the recent evidences of TEB-induced heart toxicity, the knowledge regarding the cardiotoxic effects of this fungicide in mammals is limited. Particularly, there is no functional study in the literature that describes if acute and/or long-term exposure to TEB can induce direct or indirect effects on the electrical-contractile properties of the heart and isolated cardiomyocytes. Therefore, the present investigation was conducted to uncover the effects of acute exposure to TEB on the heart excitability of adult male mice.

## Methods

### Ethical approval

All procedures involving animals were approved by the Ethical Committee on Animal Research of the Federal University of the São Paulo (protocol #1978040520), conformed to the Guide for the Care and Use of Laboratory Animals. The experiments were performed using 8 – 10 weeks old male C57BL6 mice. During procedures, all steps were taken to minimize animal pain and suffering. The mice were housed in standard rodent cages, under a controlled 12-h light/12-h dark cycle at room temperature (23 ± 2°C).

### TEB effects in L-type calcium (I_Ca,L_), potassium (I_K_), and sodium calcium exchanger (I_NCX_) currents in isolated ventricular cardiomyocytes

Recordings of whole-cell voltage-clamp were obtained using an EPC-10 patch-clamp amplifier (HEKA Electronics, Rheinland-Pfalz, Germany). Ventricular cardiomyocytes were isolated by enzymatic digestion as described by (Shioya, 2007). After achieving the proper whole-cell configuration, a 3–5 min period was waited to allow the establishment of an ionic equilibrium between the pipette solution and the intracellular environment. The internal solution for I_Ca,L_ recordings was (in mM): 120 CsCl, 20 TEACl, 5 NaCl, 10 HEPES, 10 EGTA, 1 MgCl_2_ (pH 7.2 using CsOH). The external solution used was (in mM): 150 TEACl, 0.5 MgCl_2_, 1.8 CaCl_2_, 10 HEPES, and 11 glucose (pH 7.4 using TEAOH). Regarding I_K_ currents, the composition of the internal solution was (in mM): 130 K-aspartate, 20 KCl, 2 MgCl_2_, 5 NaCl, 5 EGTA, 10 HEPES, and the external solution was (in mM): 130 NMDG, 5.6 KCl, 1.8 CaCl_2_, 0.1 CdCl_2_, 0.5 MgCl_2_, 11 glucose, 10 HEPES.

A time course of I_Ca,L_ and I_K_ current peak was recorded in a concentrationdependent manner of TEB (0.1 μM – 1000 μM). During I_Ca,L_ protocol, a pre-pulse from a holding potential of −80 mV to −40 mV for 50 ms (every 10 s) was applied to inactivate the sodium and T-type Ca^2+^ channels. After, it was applied a test pulse to 0 mV during 300 ms to measure I_Ca,L_. Considering I_K_, a pre-pulse from a holding potential of −80 mV to −40 mV was applied to inactivate sodium channels, and then the membrane was depolarized to +50 mV, thus allowing the study of I_K_.

I_NCX_ were elicited at 0.1 Hz, through a ramp protocol to −120 mV (0.09 V/s) after stepping cells to +60 mV (for 50 ms) from a holding potential of −40 mV. I_NCX_ recordings were sampled at 10 kHz, and were recorded for at least 2 min until reach stable current before exposure to 30 μmol.l^-1^ TEB. Internal pipette solution contained (in mM): 20 NaCl, 1.19 MgCl_2_, 10 HEPES, 20 TEA-Cl, 33.8 CaCl2, 40 EGTA, 40 Aspartic acid (pH 7.2 using TEA-OH), and bath solution was composed of (in mM: 145 NaOH, 145 Aspartic acid, 5 HEPES, 2 MgCl_2_, 1 CaCl_2_, 2 BaCl_2_, pH 7.3 with NaOH). Ouabain (50 μM) was added to block the Na^+^/K^+^ pump and Nicardipine (2 μM) was added to block L-Type Ca^2+^ current.

### Effects of TEB on the ventricular action potential

Ventricular action potentials (AP) were evoked by applying test pulses of 1 nA (4 ms duration). AP recordings were sampled at 10 kHz, and were recorded for 2 min with internal pipette solution containing (in mM): 130 aspartic acid, 130 KOH, 20 KCl, 10 HEPES, 2 MgCl_2_, 5 NaCl (pH 7.2 using KOH), and an external Tyrode’s solution (in mM: 120 NaCl, 2.7 KCl, 0.9 MgCl_2_, 11.9 NaHCO_3_, 1.37 CaCl_2_, 5.5 glucose, 0.4 NaH_2_PO_4_, pH 7.2). After, cardiomyocytes were incubated with 30 μmol.l^-1^ of TEB.

During AP-like protocols, cells were subjected to a voltage-clamp protocol build to mimic a short (control) and prolonged AP, typical of an AP after TEB exposure. In these protocols, cells were stepped to +40 mV (in 2ms), from a holding potential of −80 mV. The difference between them was on the repolarization phase, and a series of ramp protocols were applied to replicate the typical repolarization from mean AP records, before and after TEB exposure. Stimulation frequency was set at 1 Hz. Internal and external solutions were the same as for I_Ca-L_ records. I_Ca-L_ was obtained as the nicardipine-sensitive current, after perfusion of 2 μM nicardipine.

### Effects of TEB on sarcomere shortening

Sarcomere contractions were evaluated using a high-speed NTSC camera (MyoCamCCD100V, Ionoptix, Milton, MA, USA) through a fast Fourier transform for sarcomere deconvolution-based analysis (IonWizard, Ionoptix, Milton, MA, USA). During this experiments, freshly isolated left ventricle cardiomyocytes were placed in a coverslip bathed with Tyrode solution and assembled into a chamber containing a pair of platinum electrodes from which cells were field stimulated (MyoPacer, IonOptix, Milton, MA, USA) with 4 ms duration and 60 V amplitude biphasic pulses, at a stimulation frequency of 1 Hz. Signal was collected at a 250 Hz rate. Five consecutive events of sarcomere shortening and re-lengthening were averaged for each cell analysis. The resting sarcomere length was measured using a fast Fourier transform algorithm (IonWizard, Ionoptix, Milton, MA, USA), in a relaxed state, without stimulation. All experiments were performed at room temperature (22–25 °C).

### Effects of TEB in the intracellular calcium dynamics

Global intracellular calcium transients were elicited simultaneously with sarcomere contraction experiments, using the same stimulation protocol. Cardiomyocytes were loaded with 1 μM of the dual-excitation fluorescence probe Fura2-AM (Santa Cruz, California, USA) for 20 min, at room temperature and protected from light. Excitation was performed at 340/380 nm using a high-speed shutter (Hyper-Switch, IonOptix, Milton, MA, USA) and fluorescence emission was detected using a photomultiplier tube, controlled and digitized by the fluorescence system interface (FSI700, IonOptix, Milton, MA, USA). Five consecutive events of global calcium transient were averaged for each cell analysis. Experiments were conducted at room temperature (22-25 °C).

### *Ex vivo* electrocardiographic (ECG) experiments

For ECG, 1000 I.U. heparin was injected (i.p.), and after 15 min the hearts were removed and mounted on a constant pressure aortic perfusion system (Langendorff-like technique). Then, the hearts were perfused with Krebs-Henseleit solution (containing in mM: 120 NaCl, 5.4 KCl, 1.2 MgCl_2_, 2 CaCl_2_, 2 NaH_2_PO_4_, 27 NaHCO_3_, 11 glucose; pH 7.4) that was previously filtered (0.45 μm), oxygenated (95% O_2_ + 5% CO_2_), and heated at 34 ± 0.5 °C. To record the ECG, two electrodes (Ag/AgCl/NaCl 1 M) were placed on the heart surface, the first electrode was positioned in the ventricular apex and the second one in the ventricular base, to detect and recording the macroscopic electrical signals. All parameters were measured during 10 consecutive beats in the control condition and after perfusion with 30 μmol.l^-1^ of TEB. All signals were amplified and digitalized using a sampling frequency of 1.2 Hz (ECG-PC version 2.07, Brazilian Electronic Technology, Belo Horizonte, MG, Brazil).

## Results

### TEB blocks I_Ca,L_ and I_K_ in cardiomyocytes

Recent studies indicated that *in vivo* administration of TEB induces structural alterations in murine hearts. Here we investigated if acute exposure to TEB could directly modulate the function of ion channels in isolated cardiac myocytes. For I_Ca,L_, the membrane potential was held at −80 mV, then cells were briefly stepped to −40 mV to inactivate sodium channels, and next, the membrane was depolarized to 0 mV to elicit I_Ca,L_. Fig. 1 summarizes our findings. TEB reduced the I_Ca,L_ peak amplitude in a concentration-dependent manner (Fig. 1A-B). Tracking the repetitive measurements of I_Ca,L_ over time (not shown), it was possible to detect that the maximum stable effect of TEB at each concentration was reached in the order of a minute. Besides, when cells were exposed to increasing concentrations of TEB (from 0.1 to 1.000 μmol.l^-1^), a concentration-response curve of I_Ca,L_ peak inhibition was obtained, as plotted in Fig. 1B. The TEB IC_50_ for I_Ca,L_ inhibition was 33.2±7.4 μmol.l^-1^.

**Fig. 1.**
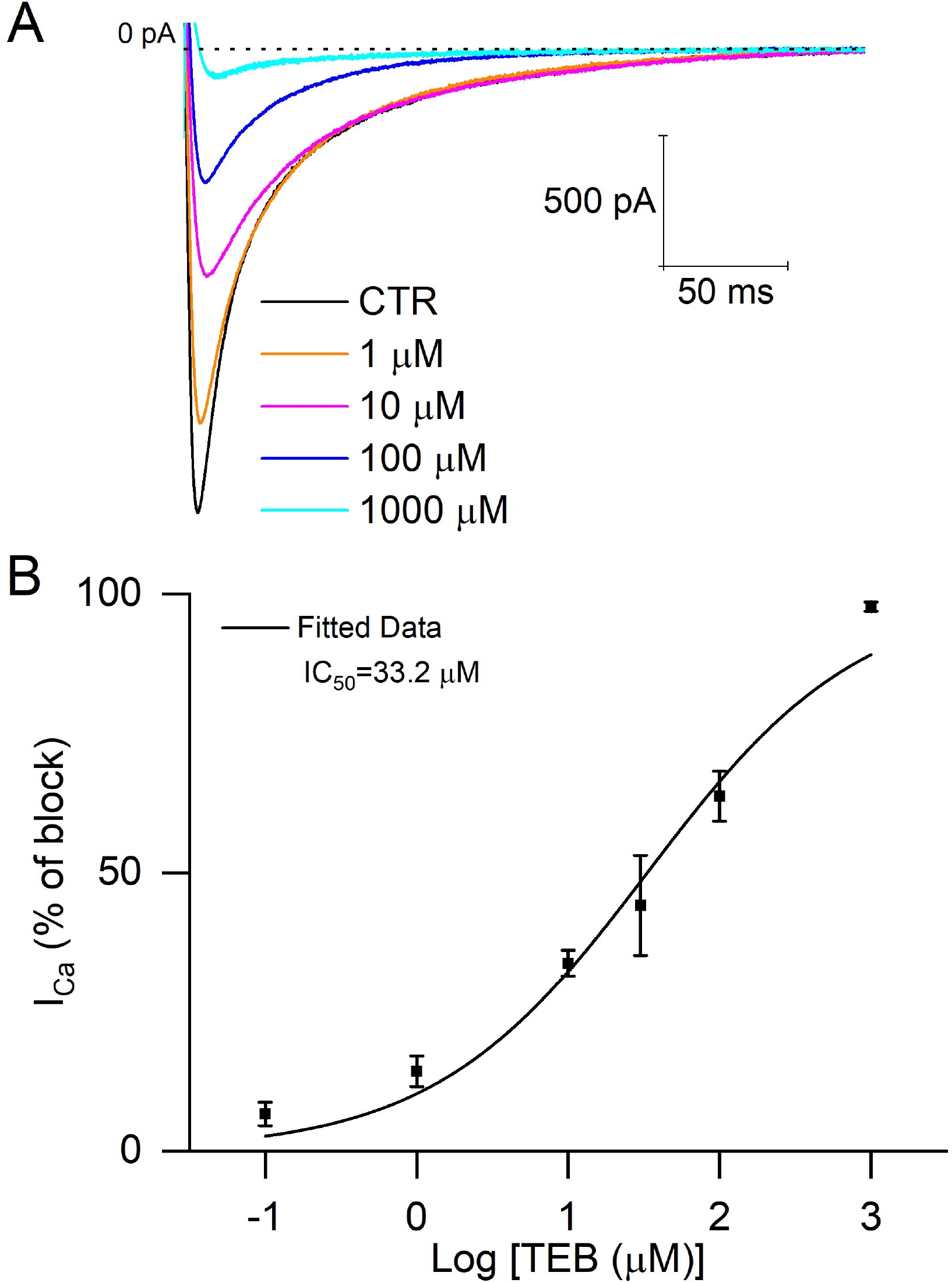
Tebuconazole blocks L-type calcium current (I_Ca,L_) in ventricular cardiomyocytes. (**A**) Representative recordings showing the I_Ca,L_ in control situation and, concentration-dependent manner of TEB (1 μM - 1000 μM). (**B**) Concentration effect curve of the I_Ca,L_ peak inhibition by TEB (IC_50_ = 33.2±7.4 μmol.l^-1^, n = 4-5 cells per point).

Since TEB was found to attenuate I_Ca,L_, we hypothesized that the pesticide could alter the function of other ion channels that control the cardiomyocytes’ excitability. To address this question, the effects of TEB on the outward potassium current (I_K_) was investigated. Starting from a holding potential of −80 mV, cardiomyocytes were stepped to −40 mV to inactivate sodium channels, and next, the membrane was depolarized to +50 mV, to elicit the I_K_. As shown in Fig. 2, TEB also was reduced I_K_ peak amplitude in a concentration-dependent manner, with ~6x higher sensibility compared to I_Ca-L_. This can also be seen in the concentration-response curve of peak I_K_ inhibition obtained when cardiomyocytes were challenged to increasing concentrations of TEB (from 0.1 to 100 μmol.l^-1^) (Fig. 2B). The TEB IC_50_ for I_K_ inhibition was 5.7±1.5 μmol.l^-1^.

**Fig. 2.**
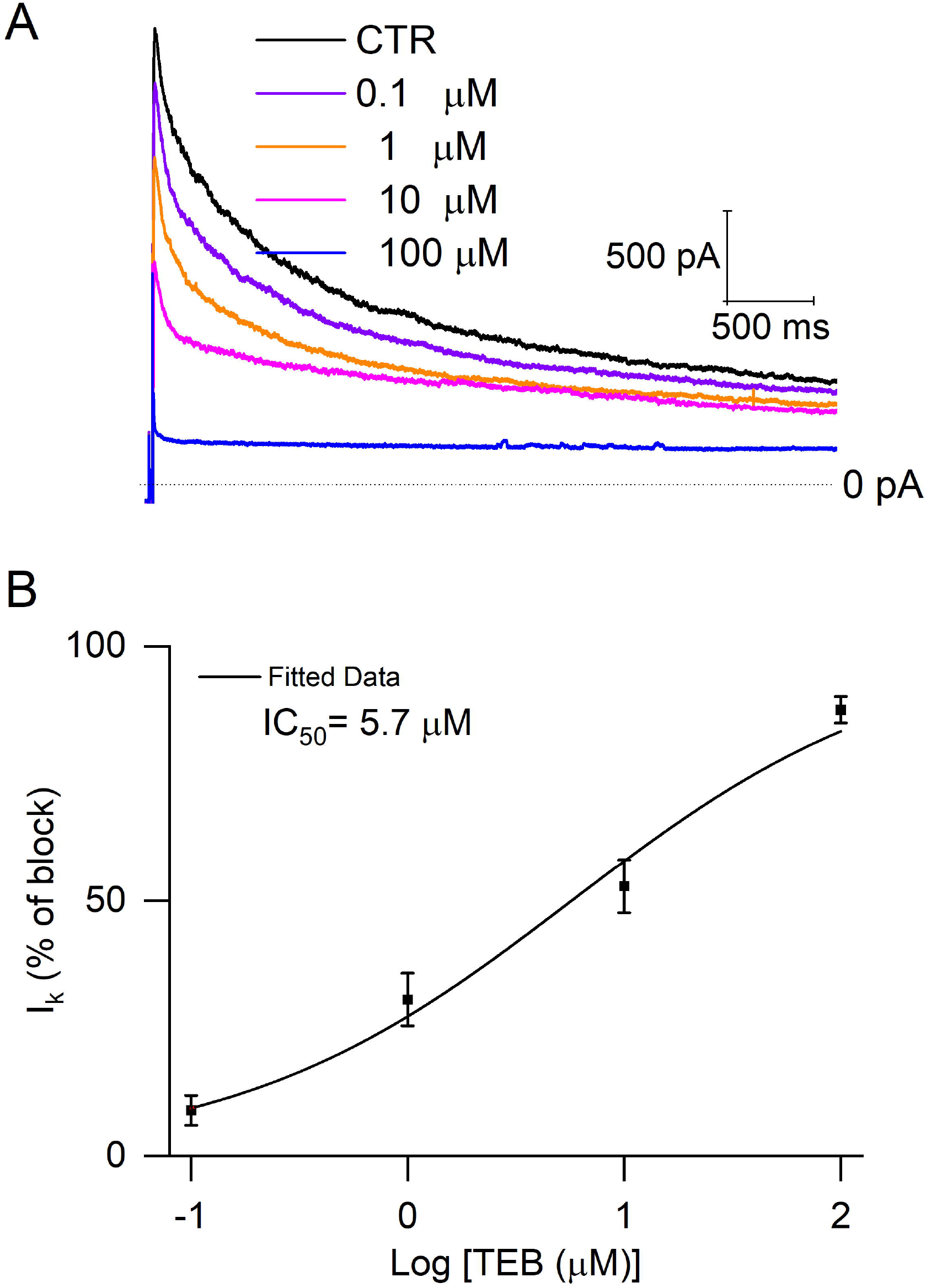
Tebuconazole blocks potassium current (I_K_) in ventricular cardiomyocytes. (**A**) Representative recording showing the I_K_ in control situation and, concentration-dependent manner of TEB (0.1 μM – 100 μM). (**B**) Concentration effect curves of the I_K_ peak inhibition by TEB (IC_50_ = 5.7±1.5 μmol.l^-1^, n = 6 cells per point).

### TEB alters the AP shape and overall excitability in isolated cardiomyocytes

To investigate TEB effects on AP parameters, cardiomyocytes were exposed to 30 μmol.l^-1^ of TEB. Concentration was chosen to trigger a significant reduction in I_Ca-L_, given its role as a central coupler of the cardiomyocytes’ excitation and contraction. Fig. 3A shows the superimposed AP waveform measured in cardiomyocytes paced at 1 Hz prior (black), during (red), and after (green) challenge cells with TEB. We evaluated time to 10, 50, and 90% of AP repolarization (APR) and it is evident that TEB was able to increase all APR evaluated (Fig. 3C, upper, middle, and down panels). Also, the effect on APR was reversible upon washout. Interestingly, TEB not only induced AP prolongation but also triggered the appearance of out-of-pace APs, an arrhythmogenic profile in cardiomyocytes (as seen in Fig. 3B, red arrow). As shown in Table 1, TEB had no significant impact on AP potential amplitude, resting membrane potential, and maximal AP depolarization rate. Thus, we conclude that 30 μmol.l^-1^ of TEB induces AP prolongation and develops arrhythmogenic phenotype in cardiomyocytes.

**Fig. 3.**
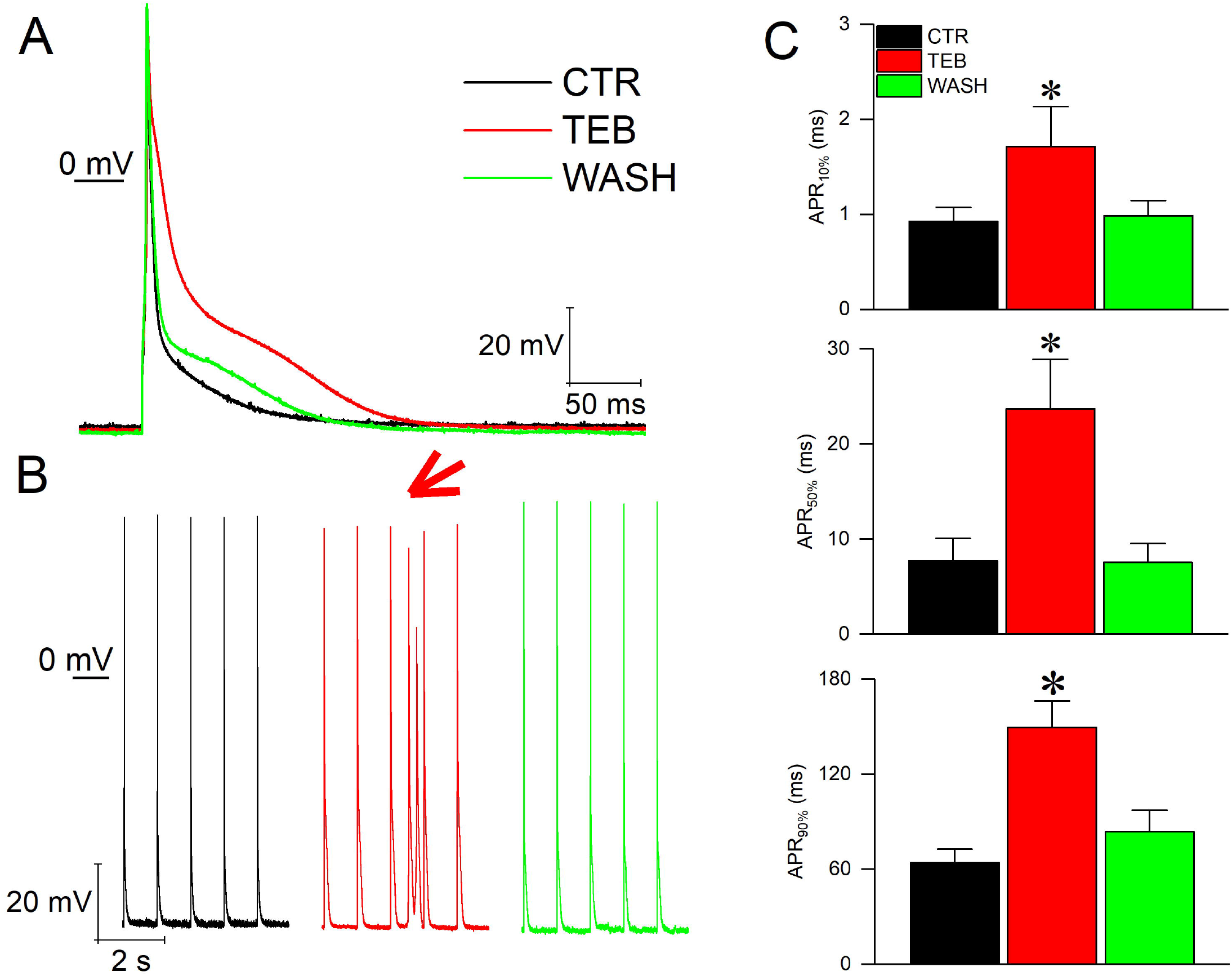
Tebuconazole changes action potential waveform in isolated cardiomyocytes and induces an arrhythmic profile. (**A - B**) Representative traces of cardiac action potential waveform (top panel) and arrhythmic profile traces (bottom) in control (black), with 30 μmol.l^-1^ TEB (red) and wash (green). (**C**) Action potential repolarization (APR) measured at 10% (top), 50% (middle) and 90% (bottom) of repolarization (n = 6 cells, *P < 0.05). Data are represented as means ± S.E.M. One-way ANOVA followed by Bonferroni post-test.

**Table 1:** Action potential parameters

|  | Amplitude (mV) | dV/dt (mV/ms) | R.M.P (mV) |
| --- | --- | --- | --- |
| **CTR** | 113.9± 3.1 | 195.4± 23.3 | -68.6± 1.2 |
| **TEB** | 112.2± 3.3 | 172.8± 19.2 | -69.7± 1.3 |
| **WASH** | 115.4±2.2 | 206.9±17.4 | -68.4± 1.7 |

### TEB increases total Ca^2+^ load during AP-simulating protocol

Despite the reduction of I_Ca-L_, our results show a significant delay of AP repolarization, given the higher sensitivity of I_K_ to TEB. Hence, we decided to investigate whether the amount of Ca^2+^ was indeed altered by the pesticide, once the repolarization of Em can impact the L-type calcium channel gating. Cardiomyocytes were stimulated with a protocol mimicking a control AP waveform, with fast repolarization (short AP). Thereafter, cells were exposed to TEB and stimulated with an AP-like protocol with slow repolarization (long AP). The AP-like protocols were created using parameters described in Fig. 4. Patch-clamp pipettes were EGTA free in these experiments, as for AP experiments. Fig. 4A displays representative traces of the I_Ca-L_ recorded from a short (left panel) and long (right panel, after a stable effect of TEB at 30 μmol.l^-1^) AP-like protocol. Solid lines are currents before exposure to 2 umol.l^-1^ nicardipine, and dashed lines are the residual current after exposure to this drug. The protocol is shown in the inset. Fig 4A, lower panels, shows the integral of the area from the nicardipine-sensitive current, used to calculate the amount of Ca^2+^ influx, since nicardipine is a selective I_Ca,L_ blocker. Nicardipine was applied in cells exposed I_Ca,L_ for both, short and long AP-like protocols, and washed out, with fully reversible effect. Fig 4B summarizes the data from the short and long AP-like protocols. Cells were paced at 1 Hz, to ensure the same frequency applied for AP experiments. The results shown in Fig. 4A (lower panel) indicate that despite the peak of inward I_Ca_ is bigger in the absence of TEB, the increase in AP repolarization time more than compensates for the total influx of Ca^2+^, even with the partial block of I_Ca_ after exposure to TEB. Taken together, these results suggest that the presence of 30 μmol.l^-1^ of TEB perfusion leads to an increase in total amount of Ca^2+^ flux during the prolonged AP, as shown in Fig. 4B.

**Fig. 4.**
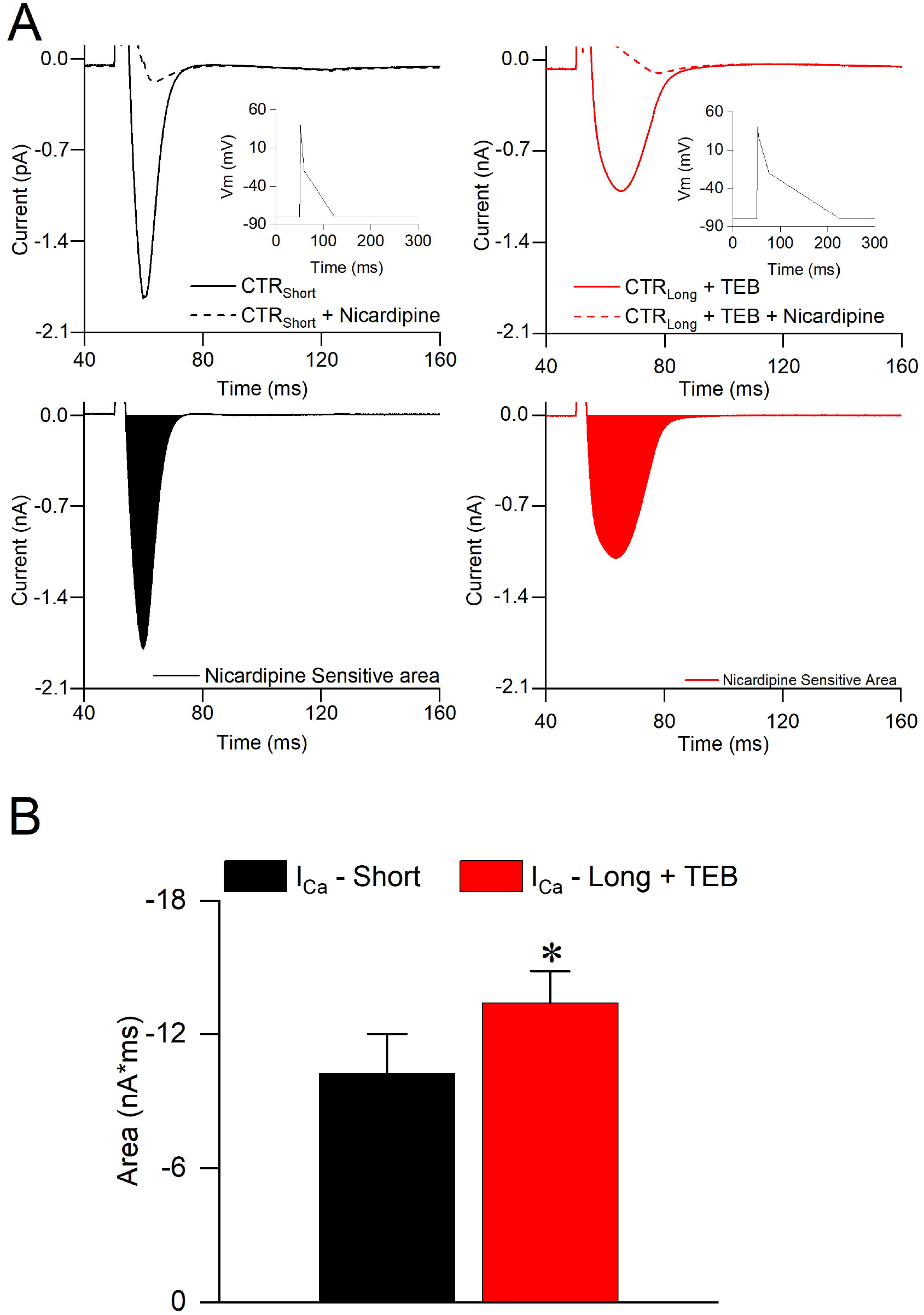
Tebuconazole increases total calcium current during action potential in cardiomyocytes. **(A)** Representative traces to mimicking AP waveform control (short action potential – left) and after challenge cells with 30 μmol.l^-1^ TEB (long action potential – right), in the absence and presence of Nicardipine 2 μM to observed I_Ca,L_ nicardipine-sensitive current. (**B**) Nicardipine sensitive area calculated as total I_Ca,L_ Nicardipine-sensitive current. Data are represented as means ± S.E.M, (n = 9 cells for each group) *P < 0.05. Student’s T-test unpaired.

### TEB enhances sarcomere shortening, calcium transient, and I_NCX_ in isolated cardiomyocytes

Given the aforementioned TEB effects on cardiomyocytes’ excitation properties, including the I_Ca-L_ and Ca^2+^ influx, we then evaluate whether cardiomyocytes contraction and calcium dynamics is further affected by this pesticide. Sarcomere shortening and Ca^2+^ transient were simultaneously tracked in isolated cardiomyocytes paced at 1 Hz, before and during the superfusion of TEB at 30 μmol.l^-1^. A significant increase in sarcomere shortening was observed (Fig. 5A, left traces), which was correlated to an increment in calcium transient amplitude (Fig. 5A, right traces). Figure 5B summarizes these findings. The contraction speed, measured as the time to reach 50% of peak sarcomere shortening was not altered (Fig. 5C, left), however, the time to 50% of peak calcium transient was delayed in the presence of TEB (Fig. 5C, right). Moreover, TEB promoted an increase in the relaxation speed, with reduced time to reach 50% of sarcomere lengthening, (Fig. 5D, left) without altering calcium reuptake (Fig. 5D, right).

**Fig. 5.**
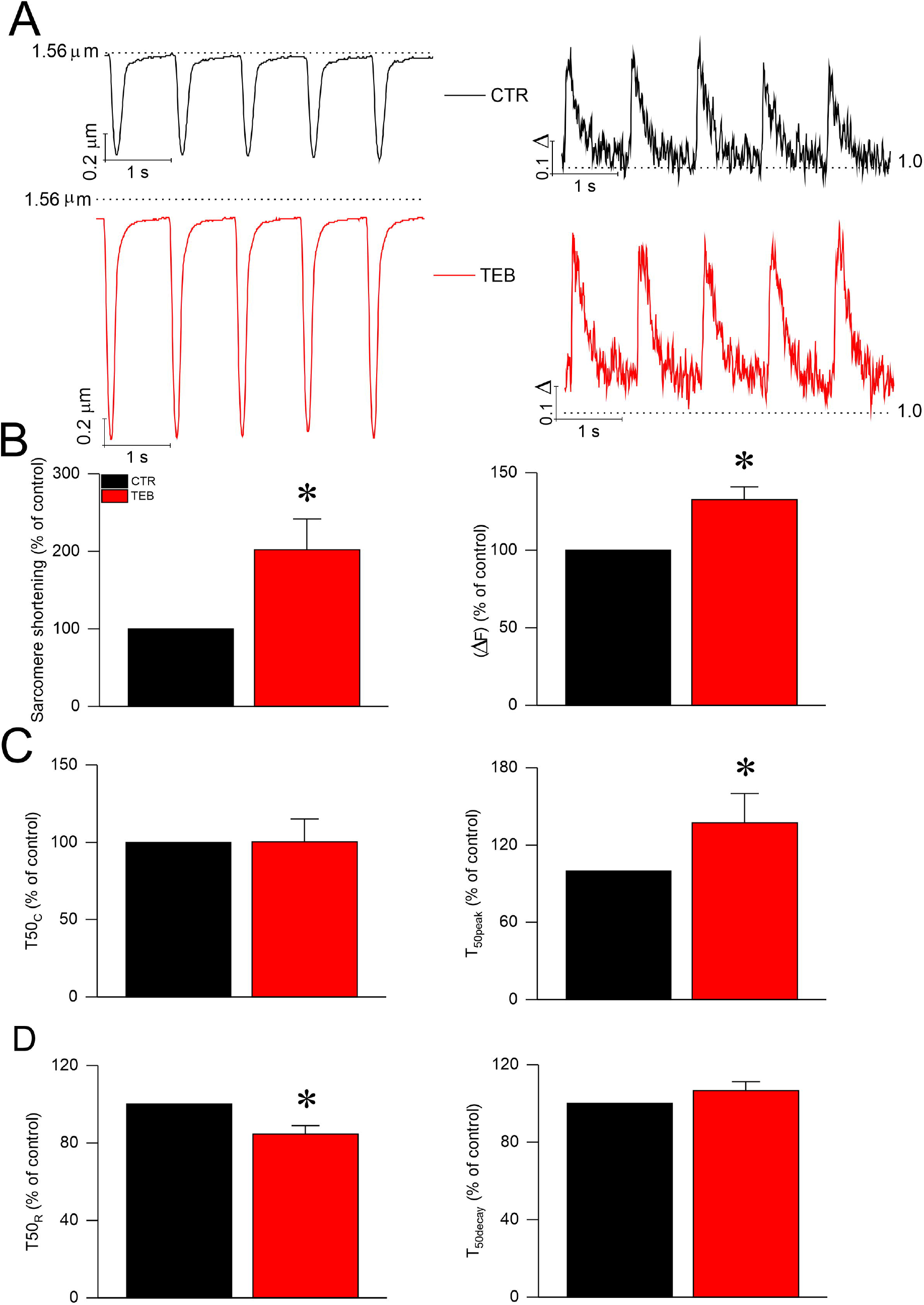
Tebuconazole enhances sarcomere shortening and calcium transient in isolated cardiomyocytes. (**A**) Five representative sarcomere contraction (left) and representative Ca^2+^ transients (right) recorded from experimental groups control (black) and presence of 30 μmol.l^-1^ of TEB (red). (**B**) Sarcomere shortening percentage (left) and global Ca^2+^ transient amplitude (right). (**C**) 50% to sarcomere shortening (left) and time to 50% of calcium transient (right). (**D**) 50% to sarcomere relaxation (left) and time to 50% of calcium decay (right). Data are represented as means ± S.E.M, (n = 10 cells) *P < 0.05. Student’s t-test paired.

As TEB appears to modulate cardiomyocyte’s contraction by altering intracellular calcium dynamics, we hypothesize that, in addition to its effects on I_Ca,L_, the pesticide may also alter the functioning of the sodium-calcium exchanger current (I_NCX_), another important sarcolemma component involved with the maintenance of intracellular calcium homeostasis. Fig. 6A provides representative records of I_NCX_ measured as a ramp protocol (inset Fig. 6A) in isolated cardiomyocytes before and after 30 μmol.l^-1^ TEB perfusion. It is possible to see that TEB increased the inward and outward components of I_NCX_ in isolated cardiomyocytes (Fig. 6B).

**Fig. 6.**
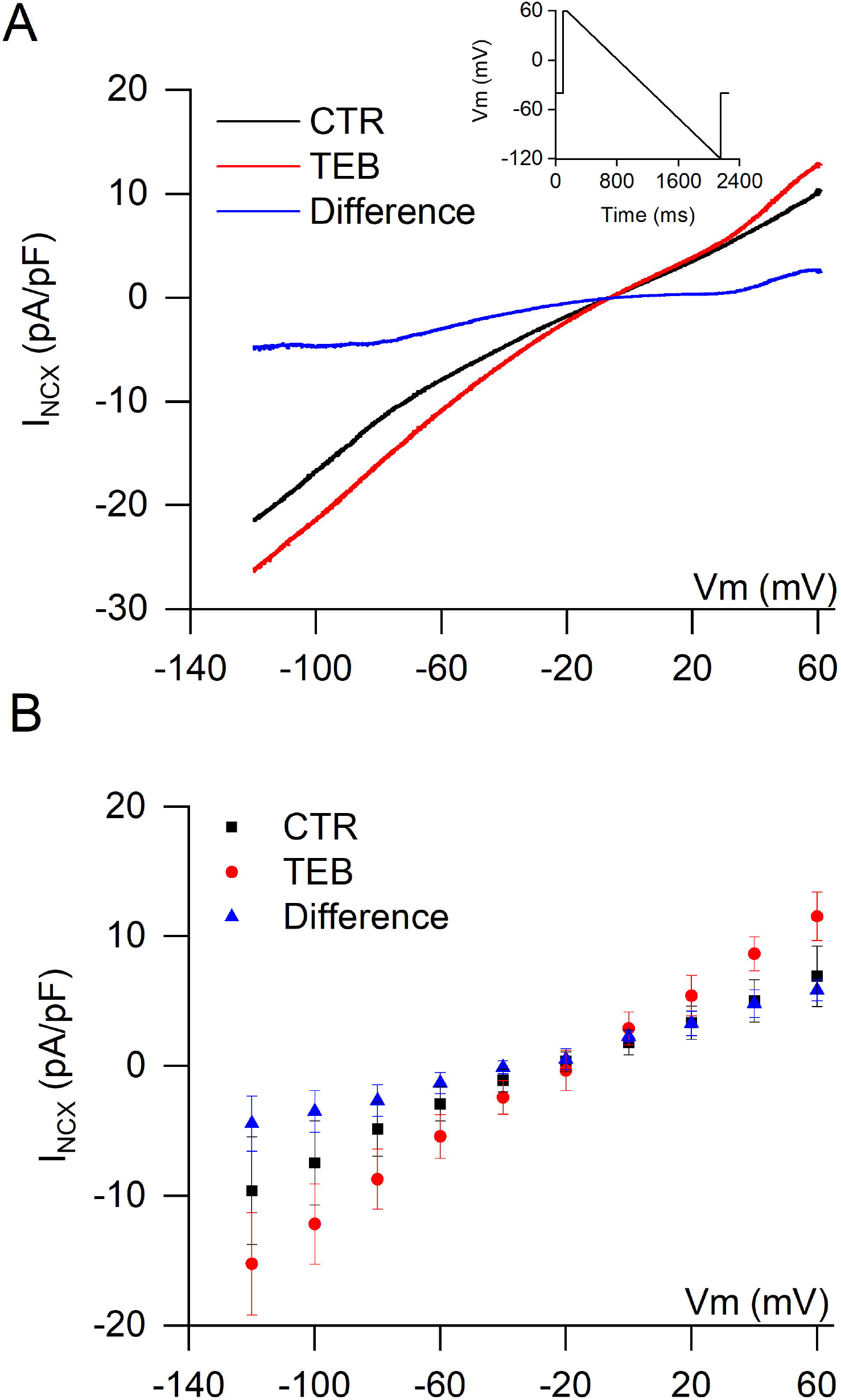
Tebuconazole increases the inward and outward component of sodium calcium exchanger current (I_NCX_) in isolated cardiomyocytes. (**A**) Representative I_NCX_ control (black) with 30 μmol.l^-1^ of TEB (red), and difference current obtained by digitally subtracting traces of control and after TEB (blue). (**B**) Average current density versus membrane potential in the control (black), 30 μmol.l^-1^ of TEB (red) and difference current obtained subtracting traces of control and after TEB (blue). Data are represented as means ± S.E.M, (n = 6 cells) *P < 0.05. Data were analyzed using Two-way ANOVA.

### TEB changes electrocardiogram (ECG) parameters and induces arrhythmias in *ex vivo* experiments

To further determine the ability of TEB to interact with cardiomyocytes and change heart electrical function, we evaluated the effect of TEB in *ex vivo* ECGs of the isolated hearts. Fig. 7A displays representative ECG traces before (black) and after 30 μmol.l^-1^ TEB perfusion (red traces). We observed that TEB induced arrhythmic phenotype in the heart. The major arrhythmic manifestation observed was atrioventricular block, as shown by the representative traces, and quantified in Fig. 7B-C. The fungicide also altered several ECG parameters. The heart rate, for example, was reduced ~50% in the presence of TEB (Fig. 7D). Furthermore, the PR, QRS, and QT intervals were lengthened by TEB, as shown in Fig. 7E-G, respectively.

**Fig. 7.**
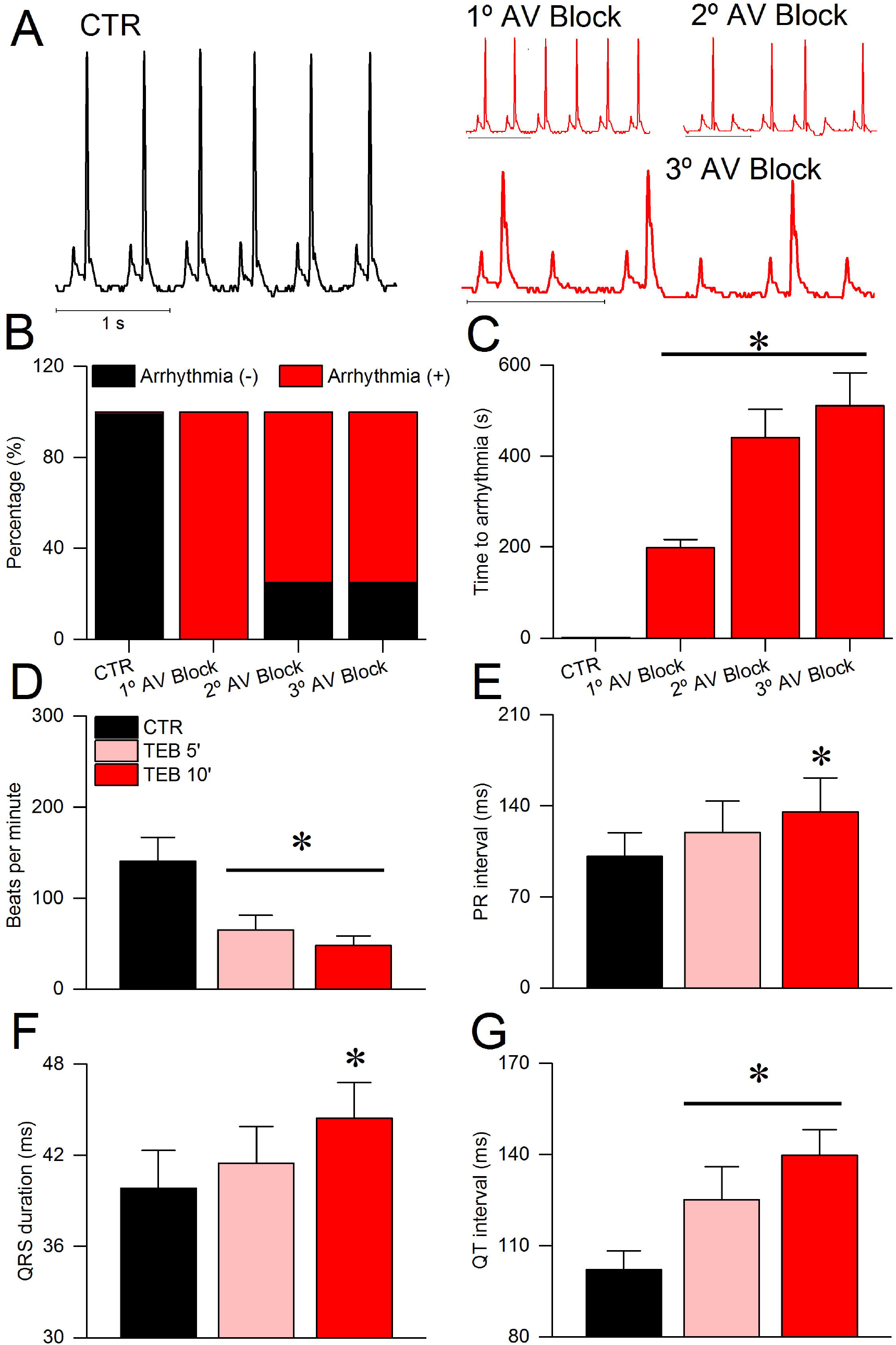
Tebuconazole changes electrocardiogram parameters and induces arrhythmias in *ex vivo* experiments. (**A**) Representative traces of electrocardiogram in control (black) and 30 μmol.l^-1^ of TEB (red) showing 1^st^, 2^nd^, and 3^rd^ AV block arrhythmias. (**B**) Percentage of occurrence of AV block arrhythmias. (**C**) Time to begin AV block arrhythmia. Myocardial electrical parameters after 5 and 10 minutes: (**D**) Heart rate (beats per minute), (**E**) PR interval (ms), (**F**) QRS complex duration (ms), (**G**) QT interval (ms). Data are represented as mean ± S.E.M. (n = 5, *P < 0.05). One-way ANOVA followed by Bonferroni post-test (**C - G**), and the Chi-square test without Yates correction (**B**).

## Discussion

In the present study, we found that TEB altered heart excitability by tracking electrophysiological features in isolated cardiomyocytes and in the ECG of isolated mice hearts. These alterations are related to a dose-dependent blockade of potassium and calcium currents and increasing in the sodium/calcium exchanger current. Modulation of ion currents by TEB led to the lengthened AP, which contributed to the occurrence of arrhythmic phenomena *in vitro*. The electrical abnormalities induced by TEB were associated with contraction remodeling in cardiomyocytes, correlated with changes in calcium dynamics. Lastly, TEB was able to promote changes in ECG parameters and also induced arrhythmias in isolated murine hearts.

Although there is evidence of the relationship between exposure to pesticides and the development of cardiovascular diseases (Berg et al., 2019; Sekhotha et al., 2016), previous toxicological evaluation reports of TEB did not describe its effects on the heart and also did not mention whether the evaluation of cardiac function was performed (EFSA, 2014; U.S. EPA, 2021). Recent studies found that TEB is indeed capable of causing remodeling of heart tissue in mammals inducing myocardial fibrosis (Ben Othmène et al., 2020b). Orally treated rats with TEB at 0.9, 9, 27, and 45 mg/kg b.w. for 28 days, showed increased levels of malondialdehyde, a marker of ROS-mediated lipid peroxidation injury, and advanced protein oxidation (Ben Othmène et al., 2020a). Various histological changes in the myocardium, including leucocytic infiltration, hemorrhage congestion of cardiac blood vessels, and cytoplasmic vacuolization were also reported. These effects of TEB were shown to be dosedependent and the deterioration of the heart was greater with the use of higher doses of TEB. The same group (Ben Othmène et al., 2020b) showed that TEB may impair parasympathetic activity in murine hearts as it reduces serum cardiac acetylcholinesterase. Interestingly, they found increased serum levels of myocardial injury biomarkers and increased levels of pro-apoptotic caspases. Additionally, TEB induced genotoxic effects by promoting DNA fragmentation and increasing the frequency of micronucleated bone marrow cells. Taken together, these results provide clues to the mechanisms by which TEB induces *in vivo* cardiotoxicity.

Regarding its biokinetic, TEB was reported to be completely absorbed in rats, goats, and laying hens treated with TEB at 2 or 20 mg/kg b.w, with a half-life in plasma varying between 31.9 and 52.5 hours. The maximum plasma concentration of TEB was approximately 4.3 μM in laying hens subjected to treatment with TEB at 10 mg/kg b.w (JMPR, 1994). Importantly, TEB accumulation in the cardiac tissue was found at an amount of 0.92 mg/kg at the end of the study. The work provided by Ben Othmène et al. (2020a, 2020b), for example, indicates that the TEB concentration range in which, supposedly, no adverse effects are observed by The no-observed-adverse-effect-level (NOAEL), can cause significant remodeling in the cardiac tissue, thus suggesting that the NOAEL should be revised.

Despite the TEB-driven remodeling of the heart and bioaccumulation in the cardiac tissue, there is currently no evaluation of the direct effects of TEB on the heart’s electrical-contractile function. Here, we found that the acute exposure of cardiomyocytes to TEB led to a significant attenuation of calcium and potassium currents. Importantly, the IC_50_ found for I_K_ was 5.7±1.5 μmol.l^-1^, almost the same value that may be found in the plasma membrane following NOAEL oral treatment for few days. Considering that I_Ca,L_ and I_NCX_ are also modulated by TEB at micromolar concentrations, this exposure may contribute to the induction of arrhythmia in cardiomyocytes and isolated hearts. Thus, based on the experimental evidence from our current investigation, we can suggest a possible mechanism involved in the induction of arrhythmias by TEB. In murine hearts, a recent study demonstrated that transient outward potassium current (Ito) is fundamental to determinate the action potential duration (APD), ECG morphology and the mechanical properties of muscle heart cells (Alarcón et al., 2018). They found that blockage of Ito led to enhanced I_Ca,L_ during AP and that I_NCX_ was also increased. Combined, these effects contributed to augmented mechanical force developed by isolated hearts and also led to prolonged QT interval (measured as QJ interval). As expected, the blocking of I_K_ by TEB induced a lengthening of the AP. The concomitant blockage of I_Ca_ by TEB would prevent such prolongation, however, TEB potency on blocking I_K_ was larger when compared with I_Ca,L_, making the AP prolongation effect dominant. In the study by Alarcón and colleagues (2018) they did not observe arrhythmic effects in the isolated heart indicating that the induction of arrhythmias by TEB may have an additional mechanism involved. Indeed, we found that TEB caused enhancement in I_NCX_ in both, outward and inward directions. Since intracellular calcium was chelated during I_NCX_ measurements, it suggests that TEB direct interacts with NCX and modulates its function.

One could argue that the TEB-driven blockage of I_Ca,L_ would result in reduced calcium transient and sarcomere shortening in cardiomyocytes. However, using AP-like protocols to mimic the control AP (short) and prolonged AP after TEB exposure (long) revealed that the blockage of I_Ca-L_ by the pesticide is more than compensated by the delay in AP repolarization promoted by attenuation of I_K_. The net result of these components is an increased total Ca^2+^ influx, since a delay in repolarization impairs the voltage-dependent component of the L-type Ca^2+^ channels inactivation. Increased calcium influx would therefore explain the subsequent increase in sarcomere contraction and Ca^2+^ transient. However, the pesticide-driven reduction of peak I_Ca-L_ may contribute to the Ca^2+^-transient kinetics, which could explain the attenuation if the time to reach 50% of total transient amplitude observed.

Nonetheless, the increase in total Ca^2+^ influx is not arrhythmogenic *per se*, which indicates that an additional mechanism is involved. TEB was reported to accumulate in the heart tissue (JMPR, 1994). Besides, it has a high partition coefficient between non-polar (octane) and polar (water) solvents (Lewis et al., 2016). Hence, it is reasonable to speculate that this pesticide may flow into the cardiomyocyte and modulate proteins found in the intracellular milieu, such as the Ryanodine receptors (RyR), although this hypothesis remains to be addressed. Importantly, here we show for the first time that TEB also enhanced I_NCX_. As demonstrated in our result (Fig. 6), I_NCX_ promotes an inward current between −45 mV and −80 mV. In its forward operation, NCX could provide a depolarizing vector during the final phase of repolarization, which may contribute to arrhythmias found during AP experiments. Indeed, arrhythmic events found in cardiomyocytes treated with TEB were trigged in a membrane potential close to −45 mV and −65 mV, which argues in favor of NCX driving arrhythmias after challenging cardiomyocytes with TEB. In some experimental models, NCX may be found as an arrhythmic substrate in the presence (Santos-Miranda et al., 2021) or absence of calcium overload in the sarcoplasmic reticulum (SR), being the latter in agreement with the present study (Hove-Madsen et al., 2004).

To extend our findings on the pro-arrhythmic potential of TEB, we conducted ECG analysis on isolated hearts. We observed arrhythmic events mainly related to sinoatrial and atrioventricular blocks. Since I_Ca,L_ is a pivotal player in the AP trigger in these cells (Bers, 2002), we can speculate that the I_Ca,L_ blockade induced by TEB may contribute to the reduction of cell excitability in the sinoatrial and atrioventricular nodes, which can explain increased atrioventricular block and reduced heart rate frequency in the presence of TEB.

Importantly, we did not observe significant ventricular arrhythmias as that observed in isolated cardiomyocytes experiments. A possible explanation for these differences is that the mean heart frequency obtained in isolated heart experiments in the absence of TEB (141.1±26.3, beats per minute), meanwhile the pacing frequency in isolated cardiomyocytes experiments was 1 Hz. Thus, apparently, the ability of TEB to induce arrhythmias is frequency-dependent. In fact, I_K_ (Dorian and Newman, 2000), I_Ca,L_ (Tiaho et al., 1994), and I_NCX_ (Omelchenko et al., 2005) are all frequencydependent and change their amplitude may impact repolarization reserve in cardiomyocytes and contribute to the occurrence of arrhythmias (Varró and Baczkó, 2011). Another possible explanation is that experiments with cardiomyocytes were performed at room temperature, meanwhile, ECG experiments were conducted at 37 °C, and ion channels and calcium dynamics can be impacted by temperature (Kiyosue et al., 1993). In the ECG we did observe QT prolongation after TEB challenge, which is well-aligned with AP prolongation observed in cardiomyocytes. Finally, conduction abnormalities, with the increase in mean PR and QRS duration may indicate that TEB could also interact with other ion channels such as the sodium channel or GAP junctions in cardiomyocytes.

In conclusion, acute exposure of murine hearts and cardiomyocytes to TEB at a low micromolar range caused significant electrical remodeling due to blockage I_CaL_, I_K_, and enhancement of I_NCX_. Combined, these effects led to increased sarcomere shortening and increment in calcium released from SR. *Ex vivo* experiments demonstrated that murine heart is sensitive to TEB and the fungicide is able to induce heart arrhythmia. From our present study, we can conclude that TEB exposure imposes a potential risk to the heart function of mammals.

